# *Ehrlichia chaffeensis* co-opts phagocytic hemocytes for systemic dissemination in the Lone Star tick, *Amblyomma americanum*

**DOI:** 10.1101/2023.08.17.553720

**Authors:** Abdulsalam Adegoke, Julia Hanson, Ryan Smith, Shahid Karim

**Affiliations:** School of Biological, Environmental, and Earth Sciences, The University of Southern Mississippi, Hattiesburg, MS 39406, USA; Department of Plant Pathology, Entomology, and Microbiology, Iowa State University, Ames, IA 50011, United States

**Keywords:** Amblyomma americanum, Ehrlichia chaffeensis, innate immunity, hemocytes, granulocyte, clodronate, immunofluorescence

## Abstract

Hematophagous arthropods can acquire and transmit several pathogens of medical importance. In ticks, the innate immune system is crucial in the outcome between vector-pathogen interaction and overall vector competence. However, the specific immune response(s) elicited by the immune cells known as hemocytes remains largely undefined in *Ehrlichi*a *chaffeensis* and its competent tick vector, *Amblyomma americanum*. Here, we show that granulocytes, professional phagocytic cells, are integral in eliciting immune responses against commensal and pathogen infection. The chemical depletion of granulocytes led to decreased phagocytic efficiency of tissues-associated hemocytes. We demonstrate *E. chaffeensis* can infect circulating hemocytes, and both cell-free plasma and hemocytes from *E. chaffeensis-*infected ticks can establish *Ehrlichia* infection in recipient ticks. Lastly, we provide evidence to show granulocytes play a dual role in *E. chaffeensis* infection. Depleting granulocytic hemocytes increased *Ehrlichia* load in the salivary gland and midgut tissues. In contrast, granulocyte depletion led to a reduced systemic load of *Ehrlichia*. This study has identified multiple roles for granulocytic hemocytes in the control and systemic dissemination of *E. chaffeensis* infection.

## Introduction

The immune system of hematophagous arthropods is important for mounting defense responses against invading pathogenic microbes. Ticks are obligate hematophagous arthropods constantly exposed to pathogenic and non-pathogenic microbes from the mammalian host during blood feeding. Pathogen recognition precedes the activation of cellular and humoral components of the immune system, which leads to the killing and elimination of pathogenic microbes from the system (Hillyer, 2011). Immune cells, known as hemocytes, play a central role in eliciting cellular and humoral responses, eventually leading to pathogen killing. Hemocytes contribute directly to immune response through phagocytosis, encapsulation, and nodulation. Hemocytes also contribute to the humoral response by producing antimicrobial peptides and lytic enzymes, which indirectly leads to pathogen killing (Fogaça et al., 2006; Kocan et al., 2008).

The recruitment of hemocytes to an infection site is necessary for efficient phagocytosis and subsequent pathogen clearance (Adegoke et al., 2023; Inoue et al., 2001; da Silva et al., 2006). Elegant studies in mosquitoes have demonstrated that hemocytes contribute to pathogen survival at the site of midgut infection and subsequent dissemination to other tissues upon acquiring an infected blood meal. Hemocytes have shown the capability to either promote or hinder pathogen survival, contingent upon the infection site, partly due to the specific attraction of these pathogens to host hemocytes. In mosquitoes, viruses such as Sindbis, Zika, and dengue are able to infect hemocytes (Parikh et al., 2009; Leite et al., 2021; Cheng et al., 2022). In ticks, *Anaplasma phagocytophilum* and *Rickettsia parkeri* also infect the tick’s hemocyte, with the former using hemocyte infection as a vehicle for salivary gland colonization (Liu et al., 2011; Adegoke et al., 2023). *Plasmodium* invasion, a mechanical process, involves the digestion of the midgut peritrophic membrane during midgut infection (Huber et al., 1991). The integrity of the midgut epithelium ensures commensal bacteria are protected and, thus, do not induce an immune response. However, damage to the midgut epithelial and basal lamina epithelial integrity, either by *Plasmodium* invasion or blood feeding, exposes the gut microbes to the epithelial, thus priming the immune system (Barletta et al., 2019). *Plasmodium-*induced midgut damage results in the recruitment of hemocytes, specifically granulocytes, to the midgut and the production of hemocyte-specific transcripts (Barletta et al., 2019; Rodrigues et al., 2010: Volohonsky et al., 2020). In addition, phagocytic granulocytes have been implicated in mosquito complement recognition of invading *Plasmodium* ookinetes, an interaction mediated by thioester-containing protein 1 (TEP1) recognition of the parasite surface that is impaired when phagocytes are chemically depleted or overloaded with beads (Castillo et al 2017; Kwon and Smith, 2019). Together, these studies strengthen the idea of phagocytes playing an integral role in the early recognition of ookinetes (Kwon and Smith, 2019).

Recent studies demonstrated that hemocytes recruited to the midgut during viral infection do not limit viral replication in the midgut; however, they are essential to restrict systemic dissemination (Leite et al., 2021). Despite the importance of hemocytes in an immune response against mosquito-borne pathogens, little information exists on the role of hemocytes in tick-pathogen interactions. Ticks can potentially harbor multiple pathogens at the same time. Unlike mosquitoes, immature and mature developmental stages of ticks can transmit tick-borne pathogens (TBPs), including a variety of bacterial, protozoan, and viral pathogens ingested during blood feeding on an infected mammalian host. In addition, some tick-borne pathogens can also be transovarially transmitted from the female to the eggs. Five hemocyte types are present in ticks based on morphological features, while functional studies have demonstrated phagocytic abilities in some hemocyte types (Adegoke et al., 2023). Granulocytes, predominant cell types, are phagocytic, thus acting as the first line of cellular defense following microbial infection. Tick-borne pathogens can infect hemocytes, specifically granulocytes, to facilitate dissemination to tissue infection, including the salivary gland and ovary. For example, *Anaplasma (A) phagocytophilum* can infect *Ixodes (I) scapularis* hemocytes and migrates to infect the salivary gland (Liu et al., 2011). Similarly, we recently demonstrated the ability of *Rickettsia (R) parkeri* to infect *Amblyomma (A) maculatum* granulocytes in both naturally and lab-infected ticks (Adegoke et al., 2023). However, events that preclude hemocyte infection, such as midgut replication, hemocyte recruitment, and subsequent pathogen trafficking through hemocytes during infection, remain a mystery.

In this study, we investigated the role of granulocytic hemocytes in the *E. chaffeensis* infection and dissemination in *A. americanum*. We provide evidence that granulocytes are integral to the tick immune response during *E. chaffeensis* infection. Our results suggest a dual role for phagocytic hemocytes in tissue infection and systemic dissemination, demonstrating that granulocytes are required to reduce tissue infection yet are required for systemic dissemination.

## Materials and Methods

### Tick maintenance and generation of infected ticks

Fully replete *Amblyomma americanum* nymphs were purchased from the Ecto services (Henderson, NC, USA) and maintained at 34□ and 65% RH under 14:10 h L:D photoperiod till needed. *Ehrlichia*-infected adult ticks were generated as previously described (Karim et al., 2012). Fully engorged nymphs were injected with *E. chaffeensis* (Arkansas strain) using a 32-gauge needle fitted to a Hamilton syringe (Hamilton Company, Franklin, MA, USA). All injected ticks were monitored for 25 hours at a temperature of 34□ to remove dead or non-viable nymphs. Viable nymphs were transferred into an incubator maintained at 34□ and 65% RH under 14:10 h L: D photoperiod and monitored until they molted into either adult males or females. Adult ticks were fed on sheep and removed at the partially fed phase (50-100mg). GFP-expressing *E. chaffeensis* (a generous gift from Dr. Uli Munderloh’s Laboratory, University of Minnesota) infected ticks were generated by injecting and capillary feeding unfed uninfected ticks with 10^7^ cell free-*Ehrlichia* suspended in sucrose-phosphate-glutamate (SPG) media (3.76mM potassium phosphate monobasic, 7.1 mM potassium phosphate dibasic, 4.9 mM potassium glutamate in distilled water) (Ammerman et al., 2008). Ticks were allowed to recover at 22°C and 95% relative humidity (RH), and hemocytes and tissues were collected to confirm infection.

### Hemolymph collection and hemocyte quantification

The collection of hemolymph and quantification of hemocytes follows the previous approach with modification to the collection medium (Adegoke et al., 2023). Perfused hemolymph was resuspended in Leibovitz’s L-15 Medium supplemented with 50% BSA on ice. Total hemocytes were quantified using a trypan blue exclusion method. Ten μL of perfused hemolymph was mixed with an equal amount of 0.4% trypan blue, and hemocytes were quantified in a Countess automated cell counter (Invitrogen, Thermo Fisher Scientific Waltham, MA, USA). Differential hemocyte count was estimated by placing 10 μL of hemolymph on the groove of an improved ‘Neubauer’s chamber. Hemocytes were differentiated morphologically as described previously (Adegoke et al., 2023, Fiorotti et al., 2019; Feitosa et al., 2015).

### Chemical depletion of phagocytic hemocytes

Unfed ticks were subjected to injection with 0.2 μL of clodronate (CLD) or control (LP) liposomes (Standard Macrophage Depletion Kit from Encapsula Nano Sciences LLC, Brentwood, TN, USA) as previously described (Adegoke et al., 2023). To identify an ideal concentration to deplete hemocytes with a minimal impact on tick survival; ticks were injected with different stocks or dilutions of LP (1:2, 1:5) or CLD (1:2, 1:5) in 1X PBS. Additionally, a group of ticks received injections of 1X PBS alone, serving as the injection control. The injected ticks were monitored for ten days following LP or CLD injection to assess their survival rates. Based on the results obtained from the optimal concentration estimation, a dilution of 1:5 (LP and CLD) was selected for subsequent depletion experiments.

### Hemolymph and hemocyte transfer

During the partially fed stage, hemolymph was collected by perfusion from CLD or LP treated E. chaffeensis-infected ticks. Hemolymph was centrifuged at 500 G for 5 minutes to separate the hemolymph plasma from the hemocyte component. The cell-free plasma and hemocyte components were maintained on ice until use. Cell-free plasma or hemocytes were injected into uninfected recipient ticks at a final volume of 0.5 μL and were subsequently fed on sheep until the partially fed stage. Partially fed ticks were collected, hemolymph isolated, and tissues and carcasses dissected. The effect of hemolymph transfer on total and differential hemocyte population was assessed as described in “***Hemolymph collection and hemocyte quantification***”. RNA was isolated from tissues and carcasses for *E. chaffeensis* quantification as described in “***RNA extraction, cDNA synthesis, and qRT-PCR***”.

### Quantification of hemocyte phagocytosis

We utilized *in-vivo* injection of fluorescent-conjugated carboxylated beads to assess and quantify hemocyte phagocytosis, as previously described by Adegoke et al. (2023) and Kwon and Smith (2019). To determine hemocyte phagocytosis, we injected ticks with 0.2 μL yellow green Carboxylated-Modified Microspheres (Thermo Fisher Scientific, Waltham, MA, USA) diluted in PBS into the tick hemolymph. Once the ticks underwent a four-hour recovery period at 22°C and 95% relative humidity (RH), we carefully perfused and processed the hemolymph for imaging. The hemolymph was placed on a cover slide and incubated at room temperature for an hour to promote hemocyte adherence to precisely quantifying phagocytosis. Following this, a 30-minute fixation step was undertaken using 4% paraformaldehyde (PFAand a 5-minute wash step in 1X PBS. The slides were then mounted and sealed in VECTASHIELD antifade mounting medium with DAPI (Vector Laboratories, catalog number: H-1000) for further image acquisition. Each group underwent processing of 3-5 ticks for the *in-vivo* phagocytosis assay to provide comprehensive data collection. For each tick, three different microscopic fields were carefully selected, each containing 200-500 hemocytes per field of view. This meticulous approach allowed us to determine the proportion of phagocytic hemocytes present.

Quantification of sessile phagocytes in *E. chaffeensis-infected* tick tissues followed the method described in the previous paragraph. The ticks used for this experiment were infected at the end of the nymphal blood meal as described in the “*Tick maintenance and generation of infected ticks*” section. Once the hemolymph had been perfused as described above, the ticks were injected with 4% PFA and allowed to fix for 1 hour at room temperature. The salivary glands and tissues were dissected in 0.5% Triton-X 100 for permeabilization for 30 minutes before washing in 1X PBS three times at 5 minutes each. The tissues were then mounted and sealed on a microscope slide using a VECTASHIELD antifade mounting medium with DAPI (Vector Laboratories, catalog number: H-1000) for image acquisition and analysis.

### Microbial infection and survival analysis in depleted ticks

Gram-negative and –positive bacteria were injected to access the immune response in hemocyte-depleted ticks. Overnight cultures of *Escherichia coli* (*E. coli*) DH5-alpha and *Staphylococcus aureus* (*S. aureus*) RN4220 maintained at 37°C in LB and TSA media were harvested, centrifuged, and adjusted to OD600 = 0.5 for and OD600 = 0.1 in 1X PBS respectively. *E. chaffeensis* (Arkansas strain) was recovered, and propagation was as previously described (Cheng and Ganta, 2021). Briefly, unfed ticks were injected with 0.2 μL of either CLD or LP (1:5) and kept to recover at 22°C and 95% relative humidity (RH). Forty-eight hours after depletion, surviving ticks were injected with 0.2 μL 10^7^ *E. chaffeensis, E. coli*, or *S. aureus*, using a 33G removable needle (Hamilton Company, Franklin, MA, USA). Heat-killed *S. aureus* and *E. chaffeensis*, and LPS were injected as positive controls. A total of 20 ticks were used per treatment or control group. Ticks were kept for recovery at 22°C and 95% relative humidity (RH) and monitored every 24 hours for ten-day survival.

### RNA extraction, cDNA synthesis, and qRT-PCR

The salivary gland, midgut, and carcass were isolated from hemocyte-depleted ticks and ticks that received hemolymph components from hemocyte-depleted ticks. Total RNA was extracted following the Trizol-chloroform separation and isopropanol precipitation method (TRI Reagent, Molecular Research Center, Inc. Montgomery, Cincinnati, OH, USA). RNA pellet was washed in 75% ice-cold ethanol, air dried, and resuspended in 30 μL nuclease-free water. The final RNA concentration and quality were checked on a nanodrop (Nanodrop One, Thermo Fisher Scientific, Pittsburgh, PA, USA), and RNA was stored at -80□ until needed. Complementary DNA synthesis was synthesized from 1000ng of isolated RNA as previously described (Bullard et al., 2016). An absolute quantification method was used to quantify *E. chaffeensis*. An absolute quantification approach establishing a standard curve of *E. chaffeensis* 16S rRNA to serve as the reference template for qPCR (Teymournejad et al., 2017). The relative expression of *E. chaffeensis* 16S rRNA gene was quantified in relation to *A. americanum* actin gene.

## Results

### Blood meal and pathogen infection induce hemocyte population changes

The microscopic examination of hemolymph collected from *A. americanum* ticks, co-stained with Phalloidin (green) and Hoechst 33342 (blue), revealed five distinct hemocyte populations. These populations exhibited specific characteristics, including Spherulocytes (Sp) with a small nucleus-cytoplasmic ratio and a peripherally displaced nucleus. Granulocytes (Gr) also displayed a large nucleus-cytoplasmic ratio and multiple cellular projections. Prohemocytes (Pr) were also present, appearing with a large nucleus-cytoplasmic ratio and smaller than other cell types. Furthermore, we identified oenocytoids (Oe), which exhibited a smaller nuclear-to-cytoplasmic ratio. Lastly, their pyriform shape and a centrally placed nucleus (Fig. 1A) distinguished plasmatocytes (Pl).

**Figure 1:**
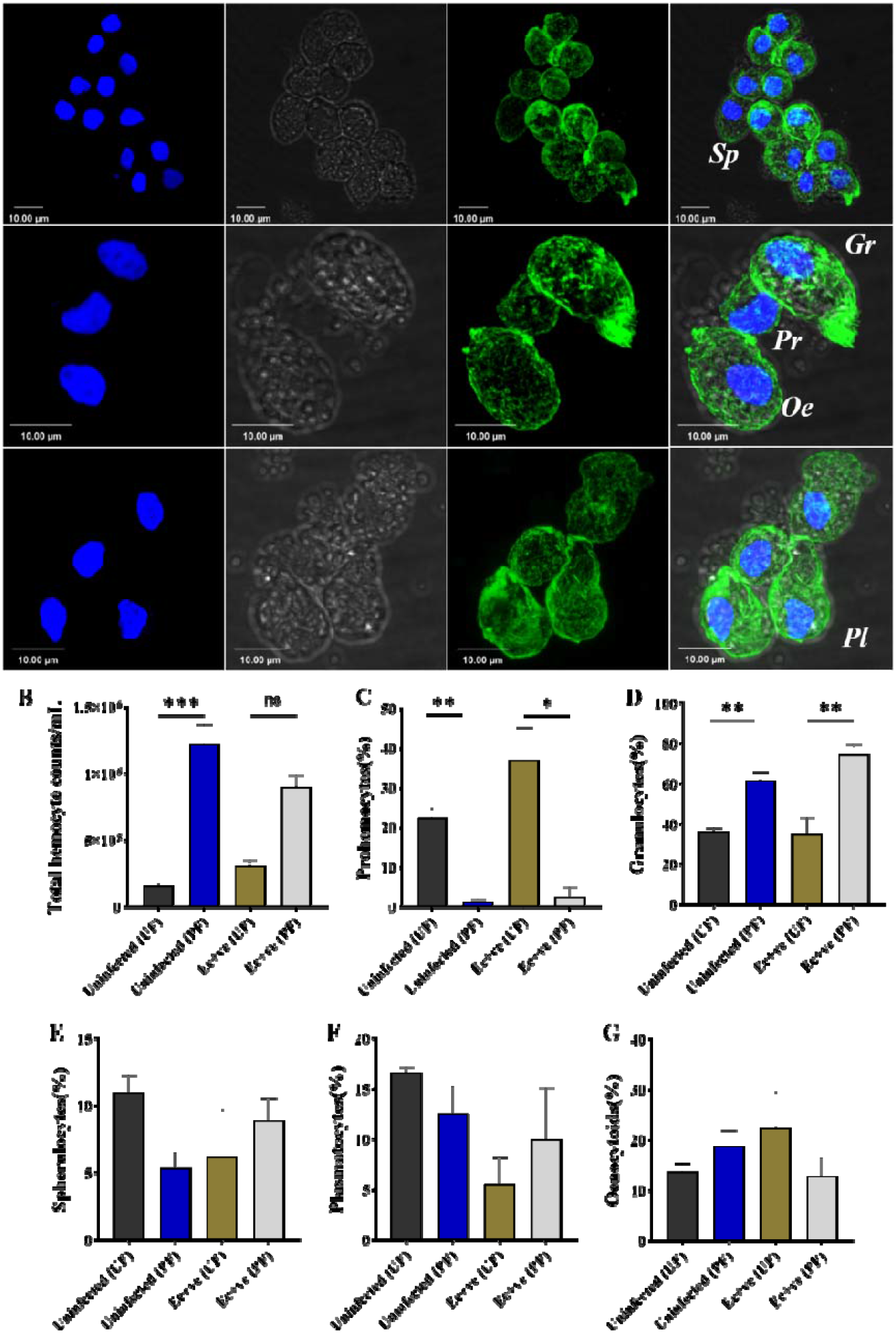
*Amblyomma americanum* hemocytes are heterogeneous with five distinct populations. Fluorescent microscopic images of *A. americanum* hemocytes stained with Phalloidin (green) and Hoechst 33342 (blue). Hemocytes are differentiated based on the size and location of the nucleus and cytoplasmic Projections. Granulocytes (Gr) displayed relatively large size and multiple actin projections. Their pyriform shape and centrally placed nucleus distinguish Plasmatocytes (Pl). Spherulocytes (Sp) possess a peripherally placed nucleus. Prohemocytes (Pr) possess a relatively large nuclear to cytoplasmic ratio. Oenocytoids (Oe) with a smaller nuclear to cytoplasmic ratio. Hemolymph was perfused from unfed and partially blood-fed (uninfected and Ec infected) ticks and the (A) total and (B-F) differential hemocyte population was compared between unfed and partially blood-fed ticks using an improved Neaubeur chamber. Data was analyzed using an unpaired t-test in GraphPad Prism 8.4.1. Asterisks denote significance (*P < 0.05, **P < 0.01, ***P < 0.001, ****P < 0.0001). UF; unfed, PF; partially blood fed. Scale bar 10 μM.

We further assess the impact of hematophagy and *Ehrlichia* infection on the hemocyte population. We counted total and differential hemocyte populations from the hemolymph of unfed and partially blood-fed ticks. Our results showed hematophagy significantly increased the hemocyte population in uninfected ticks compared to *Ehrlichia*-infected ticks (Fig. 1B). Although infection seems to stimulate a higher level of basal hemocyte counts, the effects of blood feeding are less prominent in unfed ticks (Fig. 1B). Specifically, the population of prohemocytes was considerably higher in unfed ticks compared to partially blood-fed ticks, regardless of the infection status. However, unfed *Ehrlichia*-infected ticks displayed an even higher percentage of prohemocytes when compared to unfed uninfected ticks (Fig. 1C). In contrast, we observed an opposite trend for the granulocyte population where the percentage of granulocytes were significantly higher in partially fed uninfected and *Ehrlichia*-infected ticks as compared to unfed ticks (Fig. 1D). As for spherulocytes, plasmatocytes, and oenocytoids, no significant changes were recorded following blood meal or infection (Fig. 1E-G). These results suggest that hemocytes respond to host blood meal and a lesser extent, pathogen presence through a decline in immature hemocytes and a subsequent increase in granulocyte populations. The increase in the population of granulocytes is likely in response to the overall increase in microbial load that follows a blood meal, as shown in ticks (Adegoke et al., 2023) and mosquitoes (Castillo et al., 2011; Bryant et al., 2014). Alternatively, one could posit that the physiological impacts of blood feeding also play a role, including the influence of 20E (20-hydroxyecdysone) post-blood ingestion, which has been linked to hemocyte phenotypes in mosquitoes (Reynolds et al., 2020a).

### Granulocytes are required for the immune response against commensal microbe and tick-borne Ehrlichia

Researchers have heavily relied on molecular markers to study the functions of different hemocyte types in arthropod vectors, particularly in understanding their roles in immune functions. Recent studies have demonstrated the functional characterization of phagocytic hemocytes in non-model ticks (Adegoke et al., 2023), mosquitoes, and *Drosophila* through chemical depletion to understand their roles in the immune response (Kwon and Smith, 2019; Kumar et al., 2021). Similarly in this study, clodronate (CLD) and control liposome (LP) were injected into the uninfected tick hemolymph and examined their effects on the total and differential hemocyte populations (Fig. 2A). To determine the optimal concentration for hemocyte depletion, we injected stock, 1:2, and 1:5 dilutions of CLD and LP and collected hemolymph for hemocyte quantification. Injecting CLD or LP at a 1:5 dilution had no adverse impact on tick survival (Fig. 2B). However, CLD significantly reduced the total hemocyte (Fig. 2C) and granulocyte (Fig. 2D) populations. Interestingly, CLD did not deplete the populations of prohemocytes, oenocytoids, plasmatocytes, and spherulocytes (Supplementary Fig. S1), indicating its specificity in depleting granulocytes. Assessment of hemocyte phagocytosis via *in-vivo* beads phagocytosis assay shows a significant reduction in phagocytic hemocytes as demonstrated by a decrease in the number of hemocytes associated with fluorescent beads (Fig. 2E). Next, we performed injections of CLD or LP (1:5 dilution) into *E. chaffeensis*-infected ticks and allowed them to blood feed before collecting hemolymph. The quantification of hemocytes showed a significant reduction in the total hemocyte and granulocyte population in the CLD-treated ticks (Fig. 2F-G). Additionally, *in-vivo* phagocytosis analysis confirmed a concurrent decrease in the uptake of fluorescent beads in the CLD-treated ticks (Fig. 2H-I). This result supports using clodronate liposome as an effective phagocyte depletion agent in *A. americanum* ticks.

**Figure 2:**
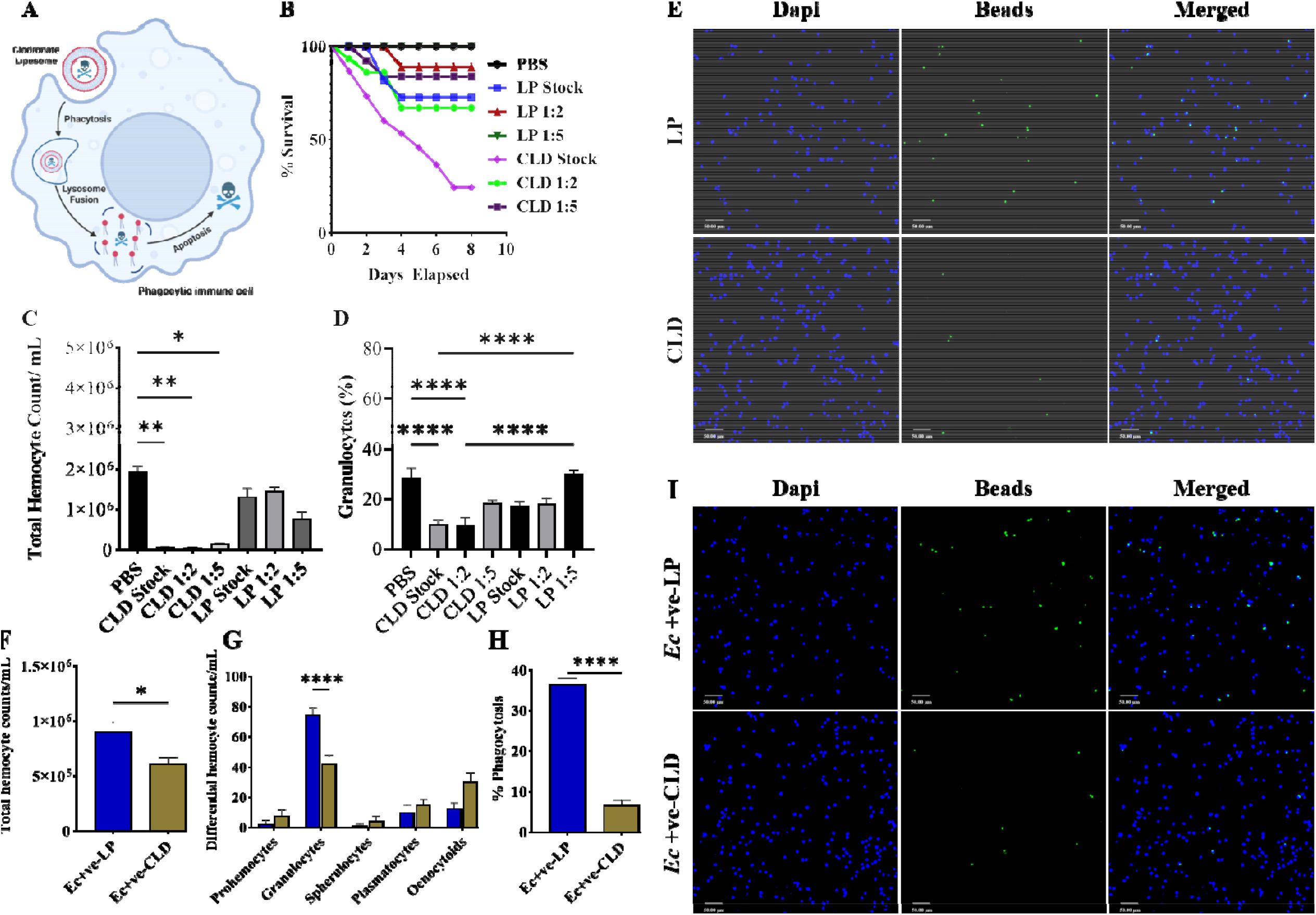
(A) Clodronate liposomes deplete phagocytic hemocyte. Different concentration of clodronate (CLD) and control liposome (LP) was injected to uninfected ticks and the hemolymph was perfused 48 hours after treatment. 1X PBS was injected as a control. (B) Ticks were monitored up to 10 days to determine the effect of CLD or LP injection on tick survival. The effect on (C) total hemocyte and (D) granulocyte population was assessed. Treatment with CLD reduced total hemocyte and granulocyte populations. (E) Hemocyte phagocytosis of fluorescent beads was also impacted in CLD treated ticks compared to control LP. Similar effect was observed when *E. chaffeensis* infected ticks were treated with CLD as shown with the reduction in (F) total hemocyte and (G) granulocyte population in *E. chaffeensis*-infected ticks. (H and I) Hemocyte depletion significantly impaired hemocyte phagocytosis of fluorescent beads in *E. chaffeensis* infected ticks. Data were analyzed using an unpaired t-test in GraphPad Prism 8.4.1. Asterisks denote significance (*P < 0.05, **P < 0.01, ***P < 0.001, ****P < 0.0001).

Ticks harbor diverse microbial communities of pathogenic and non-pathogenic commensals (Karim et al., 2021). The innate immune system actively maintains microbial homeostasis by balancing these pathogenic and commensal communities, with hemocytes playing a vital role in this process. With the goal to determine the role of the granulocyte populations in the *A. americanum* immune response, we infected ticks with *Escherichia (E*.*) coli, Staphylococcus (S*.*) aureus*, and *E. chaffeensis* after CLD treatment. As an injury control, we also included the injection of PBS. Ticks remained unaffected following the injection of PBS (Fig. 3A). CLD treatment significantly impaired tick survival following the *S. aureus* challenge (Fig. 3B). However, some ticks managed to survive until the end of the experiment. However, phagocyte depletion significantly impaired tick survival against Gram-negative *E. coli* and *E. chaffeensis* (Fig. 3C-D), with all ticks dying by the 6th and 8th day post-infection (dpi), respectively. These results underlie the integral role of granulocytic hemocytes as the first line of cellular defense during microbial infection. In addition, it demonstrates the lethality of Gram-negative bacteria in ticks when granulocytic hemocytes are significantly impaired.

**Fig. 3.**
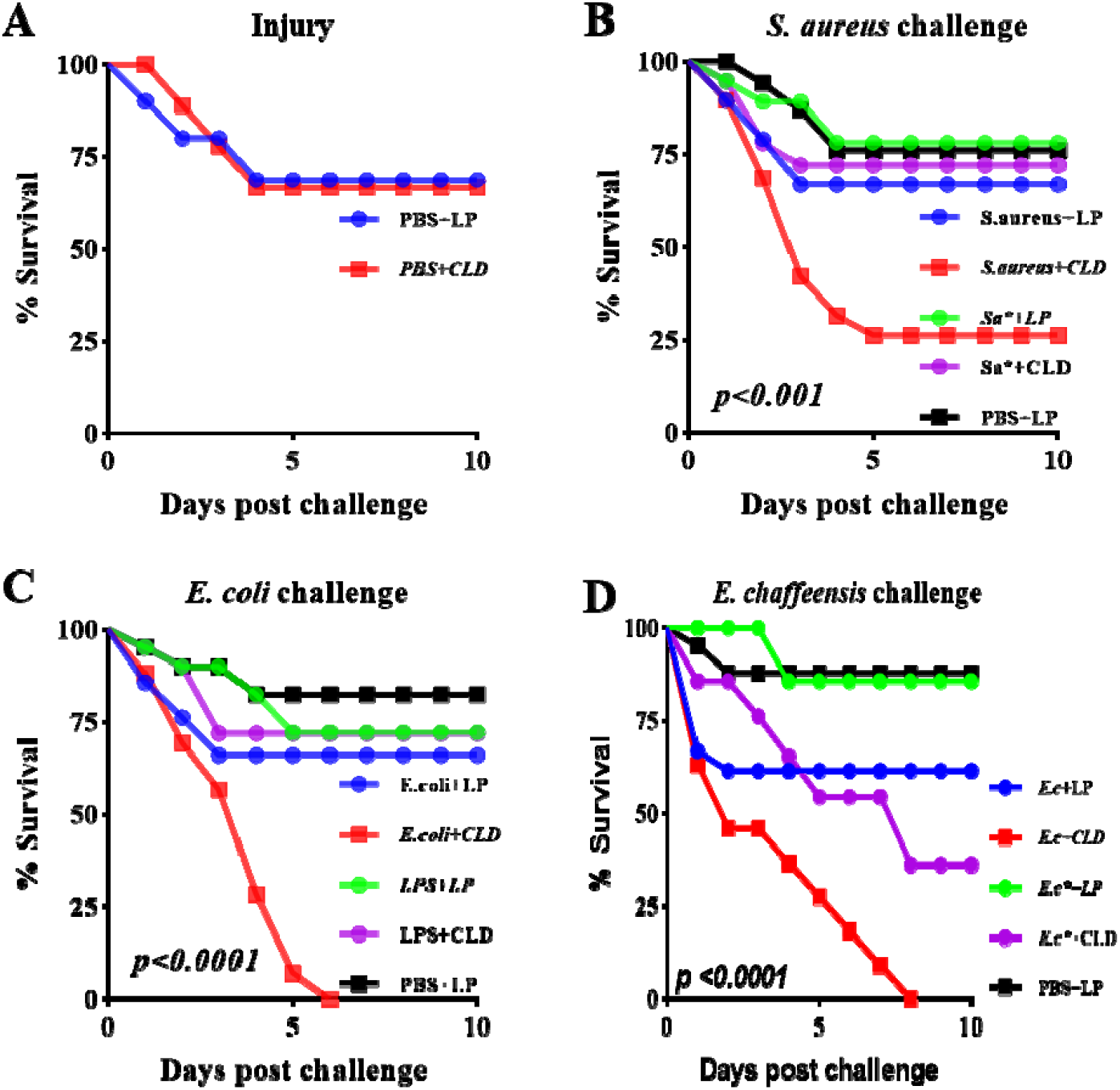
Depletion of phagocytic hemocytes impairs survival against *E. chaffeensis* infection. Ticks were injected with either LP or CLD and 24 hours after were either (A) injured or challenged with (B) Escherichia coli (*E. coli*), (C) Staphylococcus aureus (*S. aureus*) or (D) Ehrlichia chaffeensis (*E. chaffeensis*). Tick survival was monitored every 24 hours for a period of 10 days. Sterile injury (PBS injection) caused no significant loss in tick survival between CLD and LP injected ticks. Exposure of tics to Gram-negative E. coli or *E. chaffeensis* following CLD treatment led to 100% mortality before the end of the experiment. Data were analyzed by a log-rank (Mantel-Cox) in GraphPad Prism 8.4.1. LPS: lipopolysaccharide, S.a: live S. aureus, S.a*: heat-killed S. aureus, E.c*: heat-killed *E. chaffeensis*

### Phagocytic hemocytes are required to control Ehrlichia infection in tissues and are essential for systemic dissemination

Invertebrate hemocytes are distributed into two populations: freely circulating hemocytes found in the hemolymph and sessile populations attached to tissues and the body wall (King and Hillyer, 2013; Leitão et al., 2015). Several studies have demonstrated that sessile hemocytes play a crucial role in controlling microbial infection in the hemolymph from orally acquired pathogens originating from the midgut epithelium as pathogens disseminate to the hemocoel and other associated tissues (Liu et al., 2011; Rodrigues et al., 2010: King and Hillyer, 2013). To this end, we assessed the effect of clodronate treatment on the number of sessile hemocytes attached to the salivary gland and midgut of *E. chaffeensis*-infected ticks by quantifying the number of phagocytic hemocytes in these tissues. There was a significant reduction in the proportion of phagocytic hemocytes attached to both the salivary gland and midgut of CLD-treated ticks compared to the LP-injected ticks (Fig. 4A-B). In a parallel experiment, we isolated the salivary gland and midgut from *E. chaffeensis*-infected ticks 48 hours after CLD or LP treatment. Upon quantifying the *E. chaffeensis* load, no distinction was observed between the CLD and LP treated tissues (Supplementary Fig. S2).

**Figure 4:**
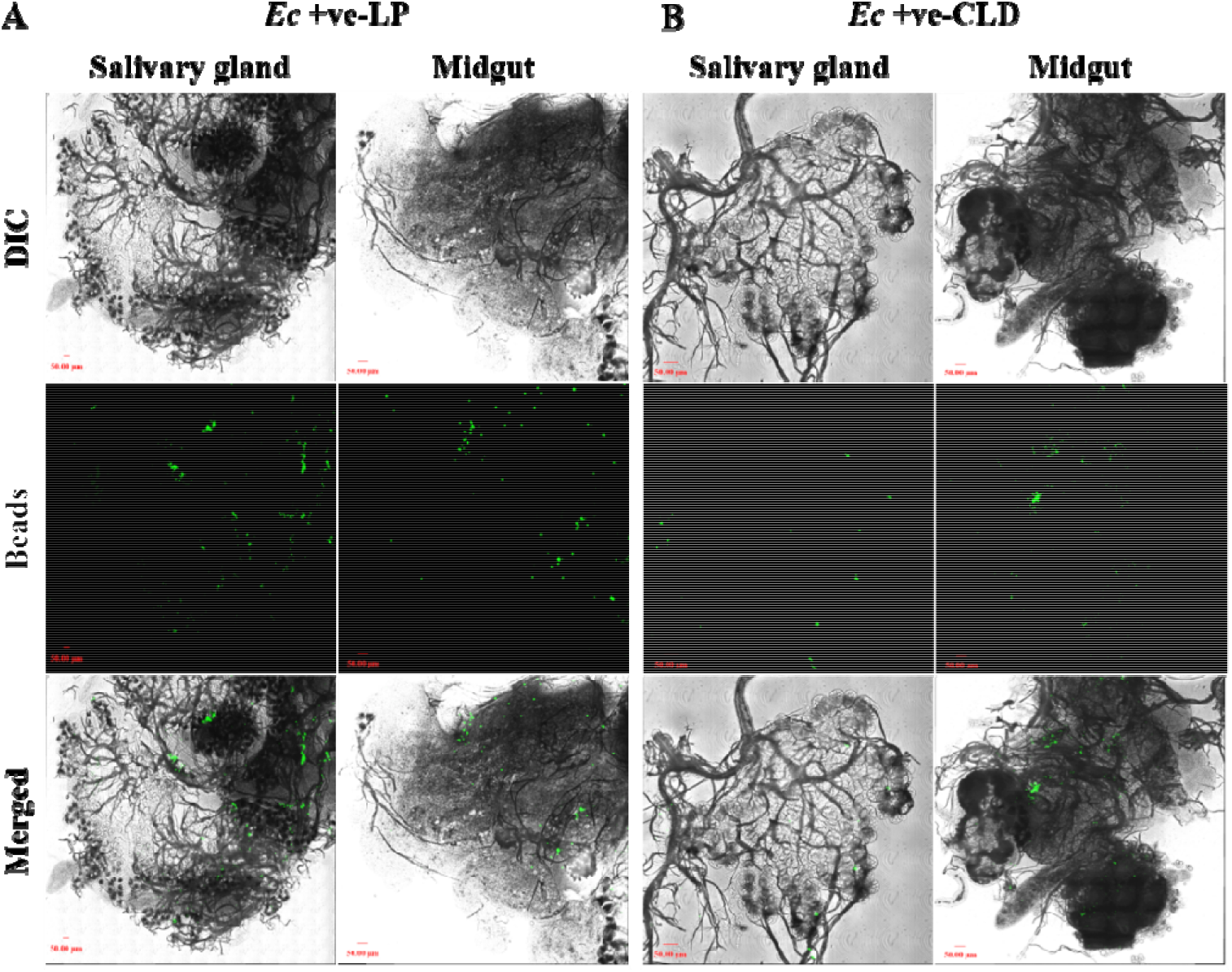
Clodronate liposome reduces the proportion of tissue-associated phagocytic hemocytes in *E. chaffeensis*-infected ticks. (A) Quantification of in-vivo phagocytosis in sessile hemocytes showed reduction in bead uptake in salivary gland and midgut from CLD treated ticks compared to LP control. (B) Hemocyte depletion has no effect on the *E. chaffeensis* load in both salivary gland and midgut compared to LP treated control. Data were analyzed using an unpaired t-test in GraphPad Prism 8.4.1. Asterisks denote significance (*P < 0.05, **P < 0.01, ***P < 0.001, ****P < 0.0001).

Tissue infection and subsequent dissemination upon infection are dependent partly on the pathogen’s ability to infect and survive inside the hemocytes (Liu et al., 2011; Leite et al., 2014). To demonstrate the ability of *E. chaffeensis* to infect hemocytes, we exposed uninfected ticks to *GFP-*expressing *E. chaffeensis* through capillary feeding and microinjection and allow the ticks to recover for 24 hours. The outcome of this experiment revealed the presence of *E. chaffeensis* in the cytoplasm of hemocytes irrespective of the infection route (Fig. 5). This further confirms that *E. chaffeensis* can infect hemocytes.

**Figure 5:**
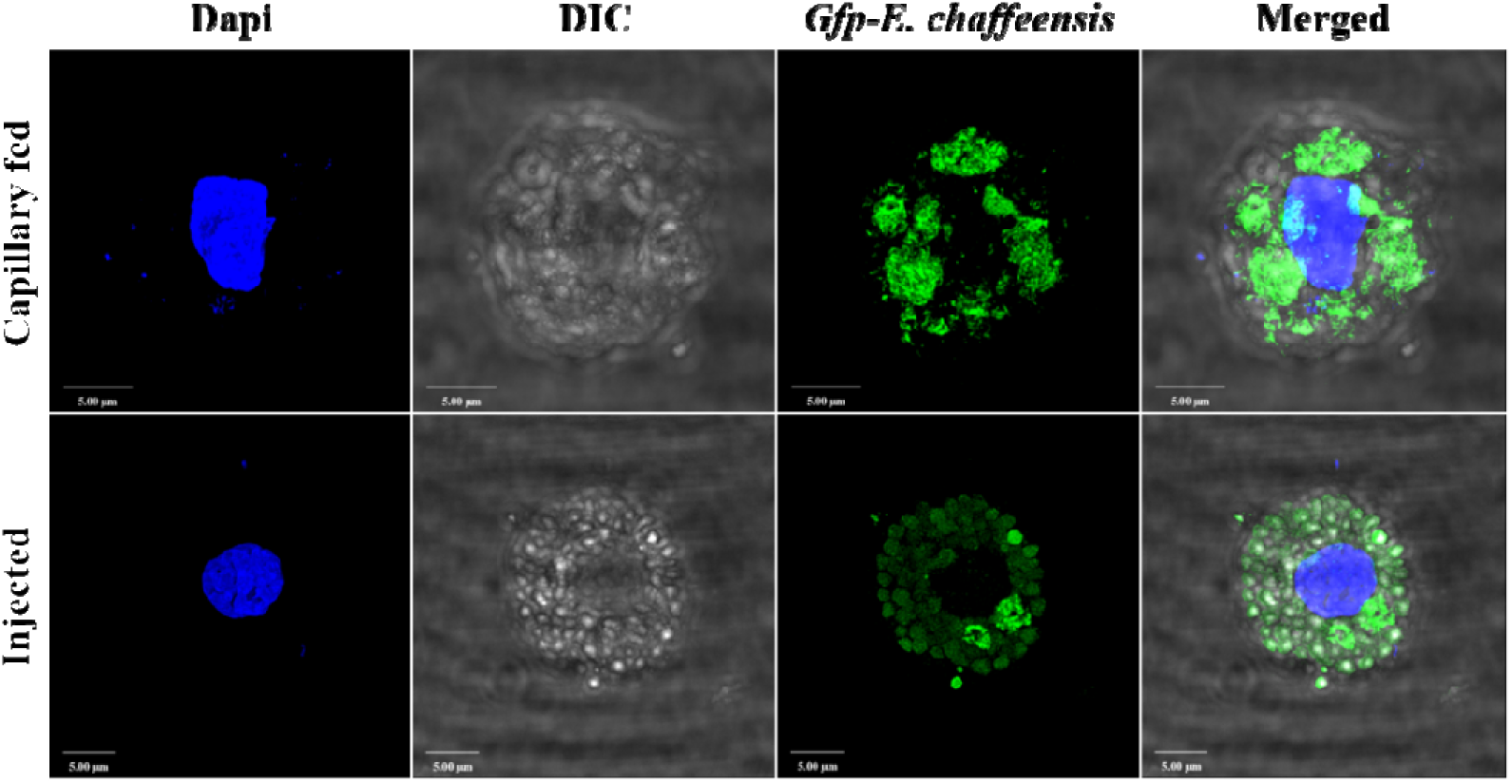
*E. chaffeensis* possess the ability to infect *A. americanum* hemocytes. Hemolymph was perfused from ticks that were previously infected with GFP-expressing *E. chaffeensis* via capillary feeding and microinjection. Immunostaining of hemocytes demonstrates the presence of intracellular *E. chaffeensis* in the hemocytes.

Here we decided to inject uninfected ticks with hemocyte or plasma (hemolymph without hemocytes) components collected from *E. chaffeensis*-infected CLD- or LP-treated ticks (Fig. 6A). In this experiment, we observed a decrease in the total number of hemocytes in ticks that received hemocyte or plasma from CLD-treated ticks (Fig. 6B). Interestingly, injection of hemocyte from CLD-treated ticks significantly increased prohemocyte and decreased granulocyte population in recipient ticks compared to recipients of plasma (Fig. 6C-D). Hemolymph components from neither CLD nor LP-treated ticks lead to changes in the plasmatocyte and oenocytoids population (Fig. 6E-F). Quantifying *E. chaffeensis* in the respective tissues and carcass of the recipient ticks, we observed varying effects of hemocyte or plasma transfer on bacterial load in tissues compared to carcass. Recipients of hemolymph components from depleted ticks exhibited a higher load of *E. chaffeensis* within the salivary gland (Fig. 6G). However, in the midgut, only individuals who received hemolymph plasma from CLD-depleted ticks displayed a higher *E. chaffeensis* load (Fig. 6G). These results in the salivary gland and tissues demonstrate that phagocytosis is essential to limit pathogen replication inside the tissues. Our results also confirmed that *E. chaffeensis* could infect *A. americanum* hemocytes as previously shown (Fig. 6G). Thus, we decided to assess the possibility of systemic dissemination via phagocytes by quantifying *E. chaffeensis* load in the carcasses of recipient ticks. Compared to the tissues, we observed a completely opposite outcome. Carcasses that had received hemolymph components from LP-treated ticks demonstrated a higher *E. chaffeensis* dissemination compared to CLD-treated ticks (Fig. 6G).

**Figure 6:**
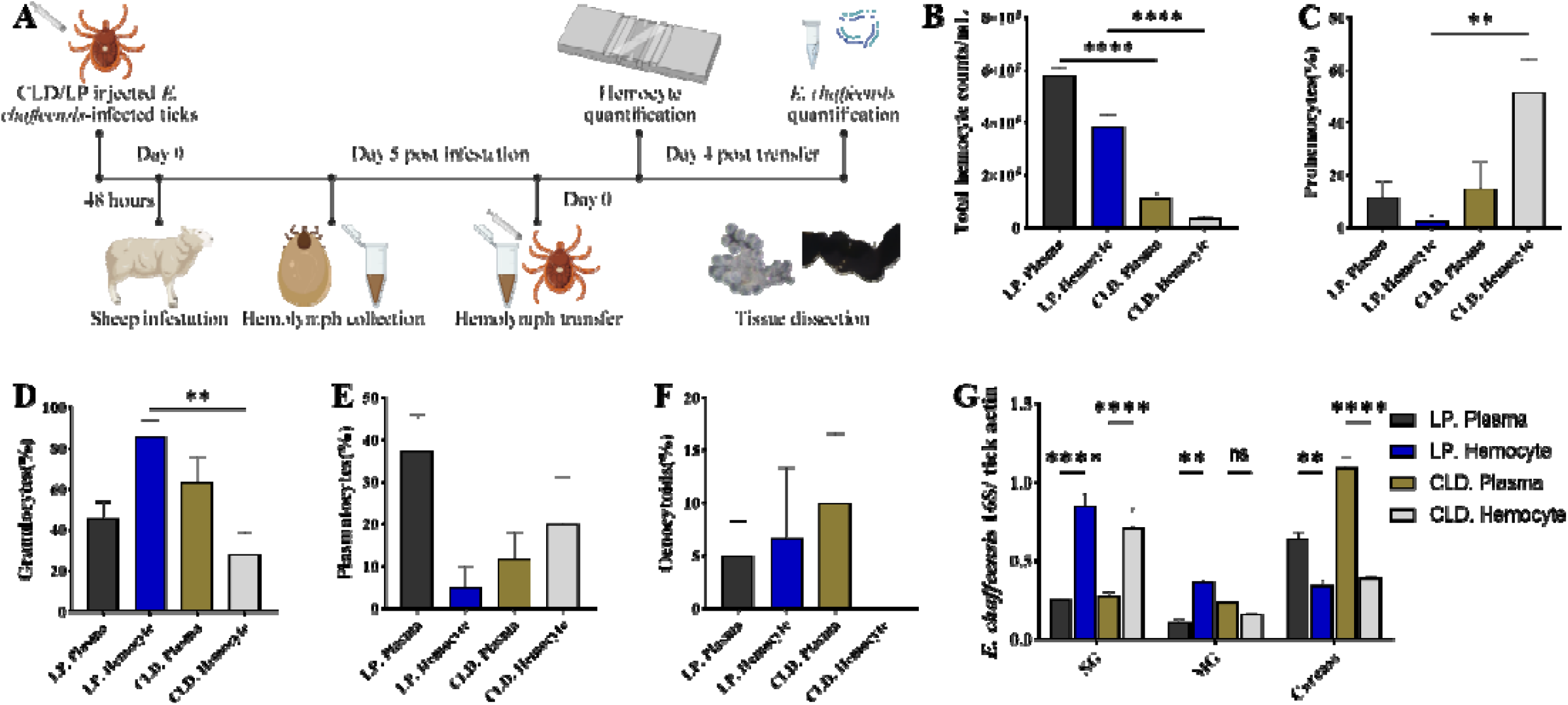
Hemolymph component from *E. chaffeensis* infected ticks establish infection in naive recipient ticks. Hemocyte and hemolymph plasma from partially fed CLD or LP treated *E. chaffeensis*-infected ticks was injected to uninfected ticks (A). The recipient ticks were subsequently blood fed prior to hemocyte and *E. chaffeensis* quantification. (BA) Total hemocyte was significantly higher in ticks that received plasma and hemocyte component from LP treated tick. Ticks that receive plasma and hemocyte component from CLD treated ticks showed a higher proportion of (CB) prohemocyte and reduced (DC) granulocyte population. No changes were observed in the population of (ED) plasmatocytes and (FE) oenocytoids. (GF) Quantification of *E. chaffeensis* in the tissues and carcass of recipient ticks showed a higher Ec load in tissues of ticks that receive hemolymph component from CLD treated ticks whereas Ec load was much higher in carcass of ticks receiving hemolymph components from LP control ticks.. Data were analyzed using an unpaired t-test in GraphPad Prism 8.4.1. Asterisks denote significance (*P < 0.05, **P < 0.01, ***P < 0.001, ****P < 0.0001).

## Discussion

In the current study, we reported the role of granulocytic hemocytes in controlling *E. chaffeensis* infection in the *A. americanum*. Previous studies from our group and others have shown that these classes of immune cells are essential for their anti-pathogen roles in non-model ticks and mosquitoes (Adegoke et al., 2023; Kwon and Smith, 2019; Leite et al 2021).

Our findings have identified five morphologically distinct hemocyte types in the hemolymph of *A. americanum*. Interestingly, the blood meal, and to a lesser extent, the infection caused by *E. chaffeensis*, contribute to changes in the hemocyte population. The hemocyte population increased following a blood meal, and *E. chaffeensis* infection, but the population of granulocytes was significantly higher in infected ticks. This suggests a heightened immune activation in response to infection since granulocytes are professional phagocytic hemocytes. Previous studies have demonstrated the use of chemical agents to deplete professional phagocytic hemocytes in *A. maculatum* (Adegoke et al., 2023), mosquitoes, and Drosophila (Kwon and Smith, 2019; Kumar et al., 2021). Partial depletion of phagocytic hemocytes significantly impairs the immune response against commensal and pathogenic microbes in invertebrates. Our study demonstrates that clodronate liposomes effectively deplete granulocyte populations in uninfected and *E. chaffeensis*-infected ticks. Likewise, we observed that ticks with depleted granulocyte populations responded poorly to microbial infection. The most intriguing finding was the complete inability of granulocyte-depleted ticks to survive *E. chaffeensis* infection. These findings support the earlier reports (Adegoke et al., 2023), where granulocyte depletion rendered *A. maculatum* susceptible to *Rickettsia parkeri* infection. These findings further strengthen the idea that granulocytes play a crucial role in mounting a cell-mediated immune response against microbial infection in the tick vector.

Due to technical difficulties with tick tissue imaging, we could not quantify the proportion of sessile hemocytes in our tick tissues. However, using *in-vivo* phagocytosis of injected fluospheres, we demonstrated that sessile hemocytes are not only present in the salivary gland and midgut tissues but are also phagocytic. The phagocytic ability of the sessile hemocytes reduced following clodronate treatment. Surprisingly, depleting sessile hemocytes in *E. chaffeensis-*infected ticks did not change *Ehrlichia* load in either the salivary gland or tissues. This could be attributed to the fact that *E. chaffeensis* was already established in these tissues before hemocyte depletion.

Infection and survival within the hemolymph and hemocyte are required for systemic pathogen dissemination in the tick vector (Liu et al., 2011). However, how these pathogens replicate and survive in the hemolymph and hemocytes is yet to be demonstrated. We observed that the transfer of hemolymph plasma and hemocyte components from hemocyte-depleted *E. chaffeensis-*infected ticks unexpectedly resulted in a drop in total hemocyte and prohemocyte population and a corresponding decline in the granulocyte population. It seems possible that the depletion in the hemocyte population is an outcome of clodronate carryover from the hemolymph plasma and hemocyte component in our transfer experiments. It is also likely that receiving hemocytes and hemolymph plasma can induce the proliferation of new hemocytes in the recipient ticks. Another important finding was that when we transferred hemolymph plasma and hemocyte components to recipient ticks; we observed changes in *E. chaffeensis* load in tissues and carcasses of recipient ticks. *Ehrlichia* infection and replication in the midgut do not rely on phagocytic granulocytes, but in the salivary gland, the presence of granulocytes is crucial to restrict *Ehrlichia* replication.. However, depletion of phagocytic hemocytes by clodronate limits systemic dissemination of *Ehrlichia*. The single most striking observation to emerge from the transfer experiment was the ability of hemocytes or plasma components from *E. chaffeensis* infected ticks to establish infection in naïve recipient ticks with varying degrees of infection. The detection of *GFP-*expressing *E. chaffeensis* in the hemocytes further suggests active replication of *Ehrlichia* in the hemolymph and hemocytes, as was previously demonstrated for *R. parkeri* (Adegoke et al., 2023) and *Anaplasma phagocytophilum* (Liu et al., 2011).

In conclusion, this study sheds light on the critical role of granulocytic hemocytes in the immune response against *E. chaffeensis* infection in *A. americanum ticks*. The presence of *E. chaffeensis* in tick hemocytes demonstrates the potential for intracellular infection and dissemination within the vector. Additionally, we identified the importance of granulocytes in limiting *E. chaffeensis* replication in the salivary gland, while their presence is not necessary for midgut infection. Furthermore, our data suggest that *E. chaffeensis* infection can alter the hemocyte population, leading to changes in the immune response of the tick. These findings highlight the complexity of vector-pathogen interactions and underscore the need for further research to elucidate the precise mechanisms by which tick hemocytes modulate immune responses against understudied tick-borne pathogens as is the case with *E. chaffeensis*.

## Data availability statement

### Ethics statement

Protocols for tick blood-feeding were approved by the University of Southern Mississippi’s Institutional Animal Care and Use Committee (USMIACUC protocols #15101501.3 and 17101206.2).

## Author contributions

Conceptualization: AA, RCS, SK; Data Curation: AA, JH, SK; Formal analysis: AA, JH, RCS, SK; Funding acquisition: SK; Investigation: AA, JH, SK; Methodology: AA, JH, RCS, SK Project Administration: SK; Resources: RCS, SK; Supervision: SK; Writing original draft: AA, JH, SK; Writing, reviewing & editing: AA, JH, RCS, SK

## Acknowledgments

We thank Dr. Uli Munderloh, University of Minnesota, for the generous gift of the GFP-expressing *Ehrlichia chaffeensis* strain, Dr. Chris Paddock, CDC, for the generous gift of *Ehrlichia chaffeensis* (Arkansas strain), and Latoyia Downs for her technical help in culturing *E. chaffeensis* strains.

