## Supplementary material for "*Ehrlichia chaffeensis* co-opts phagocytic hemocytes for systemic dissemination in the Lone Star tick, *Amblyomma americanum*": Fig S1, Fig S2

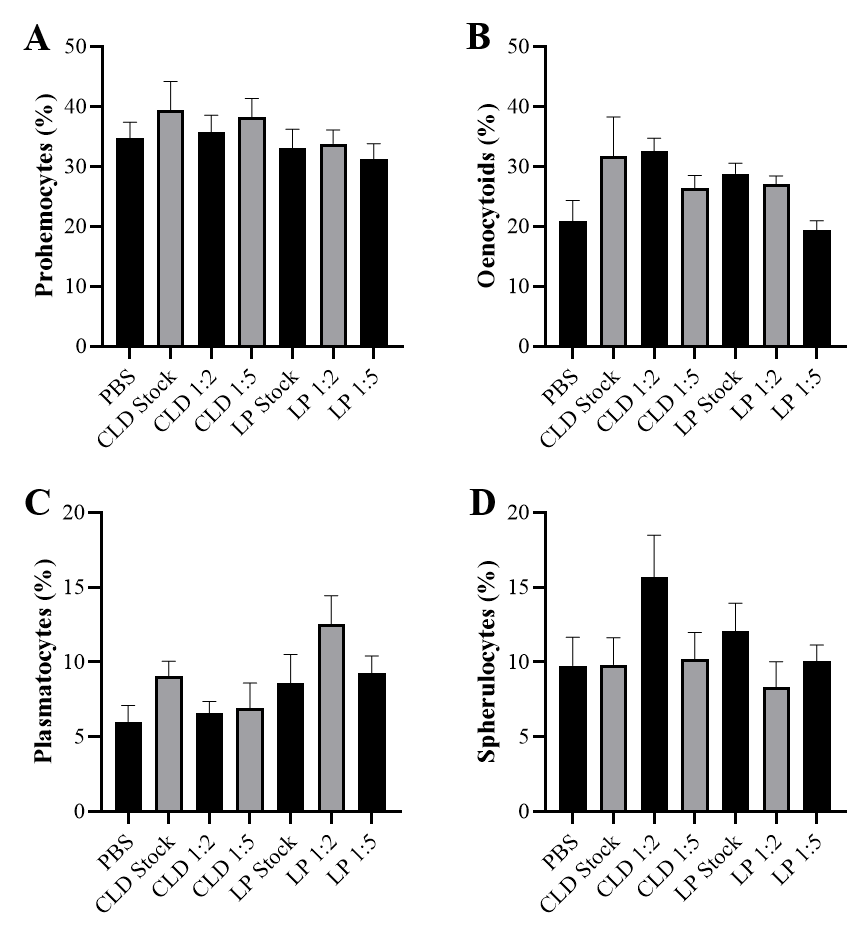
**SUPPLEMENTARY FIGURES**

Figure S1: Clodronate liposomes deplete phagocytic hemocyte. Different concentration of clodronate (CLD) and control liposome (LP) was injected in unfed ticks and the hemolymph was perfused 48 hours after treatment. 1X PBS was injected as a control. The effect on (A) total and (B-F) differential hemocyte population was assessed. Treatment with CLD reduced total hemocyte and granulocyte populations. Data were analyzed using an unpaired t-test in GraphPad Prism 8.4.1. Asterisks denote significance (*P < 0.05, **P < 0.01, ***P < 0.001, ****P < 0.0001).

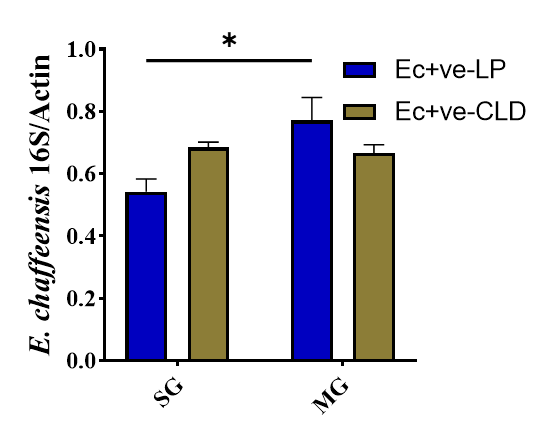

Figure S2: Hemocyte depletion has no effect on the *E. chaffeensis* load in both salivary gland and midgut compared to LP treated control. Data were analyzed using an unpaired t-test in GraphPad Prism 8.4.1. Asterisks denote significance (*P < 0.05, **P < 0.01, ***P < 0.001, ****P < 0.0001).
